# A Pilot-Scale Process for the Extraction of Raffinose-Oligosaccharides from Pulse Protein Concentrates

**DOI:** 10.1101/2024.04.19.590199

**Authors:** Philipp Garbers, Sara M. Gaber, Catrin Tyl, Stefan Sahlstrøm, Svein H. Knutsen, Bjørge Westereng

## Abstract

Plant protein concentrates have gained popularity in recent years as part of a shift towards healthier and sustainable food production. Their production via dry fractionation is well established and leads to a protein ingredient that is also rich in raffinose family oligosaccharides (RFOs). Although these can cause gut discomfort, they might also serve as a valuable substrate fowr growth of probiotic bacteria or as substrate in other biotechnological applications. By using a process including mildly acidic extraction, separation and sequential filtration, a carbohydrate fraction enriched in RFOs was isolated from peas (48 %) and faba beans (26 %) on a kilogram scale. Simultaneously a fraction with elevated protein concentration (5-6 % increase compared to the starting material) and only minor changes in amino acid composition is produced. This process could serve ingredient manufacturers that seek to reduce RFOs in protein concentrates or to produce RFOs for biotechnological or chemical conversion towards new products. It also has potential to reduce the amount of acid and base compared to protein isolate production through wet fractionation. Furthermore, the whole process was performed with industrially relevant and scalable equipment.

**Graphic for table of contents:** 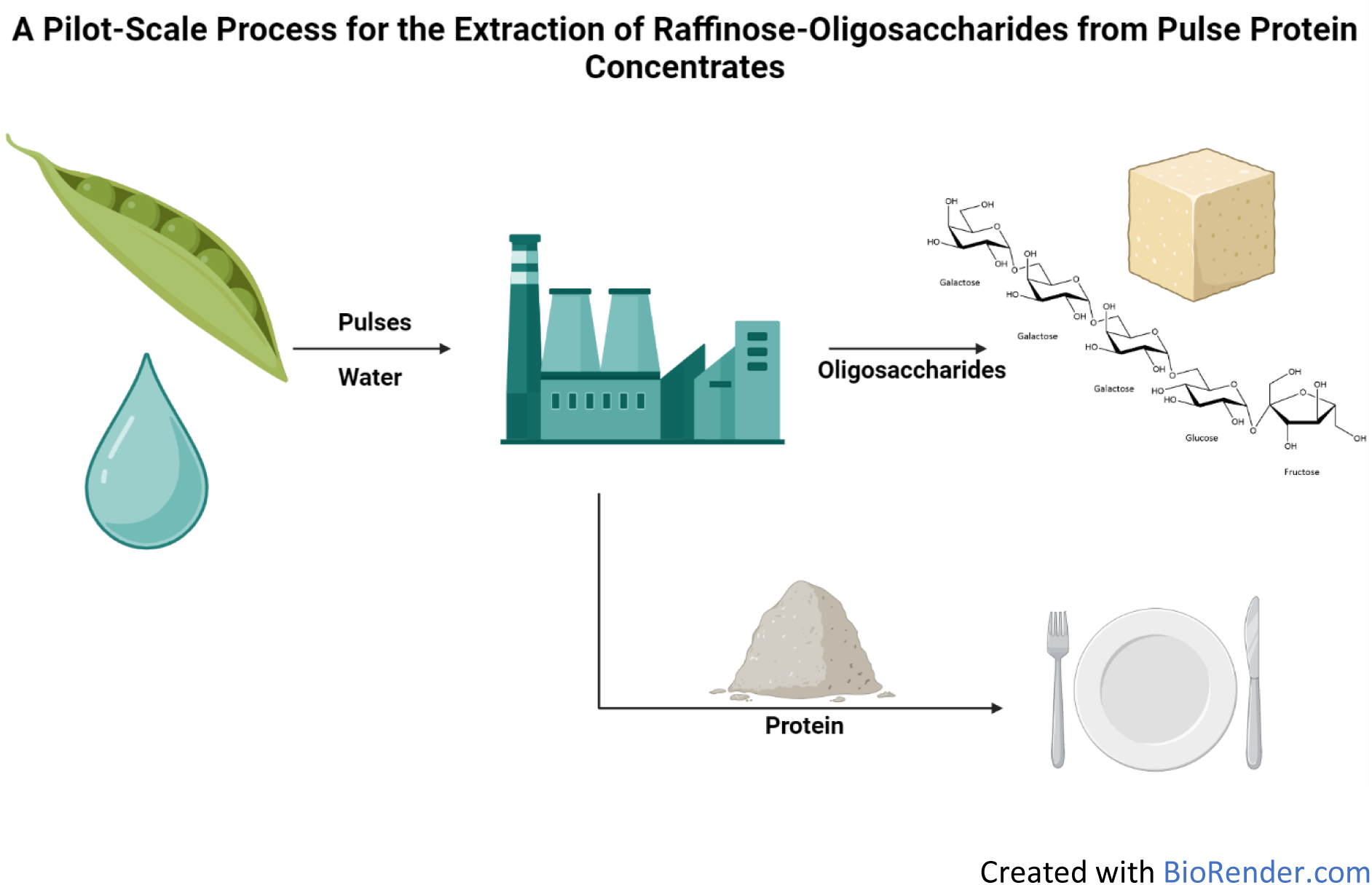

## 1. Introduction

Pulses such as field peas or faba beans have gained much attention in recent years as alternative protein sources to either animal or soy protein. Compared to soy, pea and faba bean can be grown in colder climates, for example in Norway^1^, Finland^2^ or Canada^3^. However, globally they are underutilized and grown to a much lower extent than soybeans^4^. At the same time, they convey environmental and health benefits when integrated into agriculture and diet, mainly through their protein and dietary fiber content, but also the plants’ ability to fixate nitrogen and thus reduce fertilizer use^5^. The recently published Nordic Nutrition Recommendations therefore advise for an increased consumption of pulses^6^. Pulse proteins have become especially popular in their texturized form as an ingredient in meat-analogue products^7^, but also as functional ingredients^8^. Many studies therefore focus on the functional properties of proteins from different plants and the difference between protein concentrates, isolates, and combinations thereof^9–11^. Concentrates are commonly produced by dry-fractionation^12,13^ which is considered more environmental friendly, albeit at the cost of lower protein content with higher ash, oligosaccharide (e.g., raffinose) and fiber concentrations^12,15^. Protein isolates on the other hand are produced by alkaline solubilization (e.g., via sodium hydroxide) and subsequent acidic precipitation (e.g., via sulfuric)^13,14^ and contain more protein than the concentrates but require substantially more water and chemicals and can lead to denaturation^11,12^. Some studies have also suggested combined wet and dry processes^16,17^.

Besides proteins, pulses are also rich in raffinose family oligosaccharides (RFOs). These oligosaccharides consist of sucrose that is α-1,6 linked to one to three galactose units via the C6 of the glucose end (Figure 1). The resulting trimer is called raffinose, the tetramer stachyose and the pentamer verbascose. Peas contain 2.26-6.34 %^18,19^ and faba beans between 1.40 and 3,67 %^18,20^ of RFOs. For their complete breakdown an invertase / sucrase (GH32) and an α-galactosidase (GH36) are necessary^21^. As humans lack an α-galactosidase, they cannot digest these oligosaccharides. Instead, these oligosaccharides are easily fermented by the gut bacteria in the lower intestine and are therefore considered fermentable oligo-, di- and monosaccharides and polyols (FODMAPs)^11,22,23^. The bacterial fermentation leads to gas production and can cause discomfort, bloating, nausea or even diarrhea when ingesting RFOs in larger amounts^18^, which can pose a challenge for people with gastrointestinal conditions such as irritable bowel syndrome (IBS)^21^. On the other hand, as RFOs are fermented by gut bacteria, there is an inherent prebiotic potential which has however not been sufficiently researched and only Japan recognizes RFOs as prebiotics for humans^18^ and some studies investigated RFOs as a prebiotic for animals (e.g., chickens)^24^. Furthermore, there are several examples in literature that show a potential to utilize RFOs. One approach would be to produce lactic acid through fermentation with lactic acid bacteria (LAB) as previously shown with the vinasse (leftovers) from alcoholic fermentation of soybean molasses where RFOs remain as a carbon source for LAB^25^. Fermentation could also be used to produce single cell protein from different food-grade organisms, for example the fungi *Rhizopus* (from tempeh) of which some strains can grow on RFOs as their sole carbon source^26^. This way starter cultures for tempeh or the biomass itself as an ingredient could be produced. RFOs could also be added as a selective substrate for fermentations with multiple microorganisms, where only one or multiple desired organisms can ferment them. This has previously been demonstrated with wood-derived oligosaccharides in the production of sour beer^27^. An example for a different, potentially more value-added approach, would be to use alternansucrase to produce new oligosaccharides from raffinose and sucrose which could act as prebiotics and promote beneficial gut bacteria (such as *Bifidobacterium*) without benefitting pathogens^28^.

**Figure 1:**
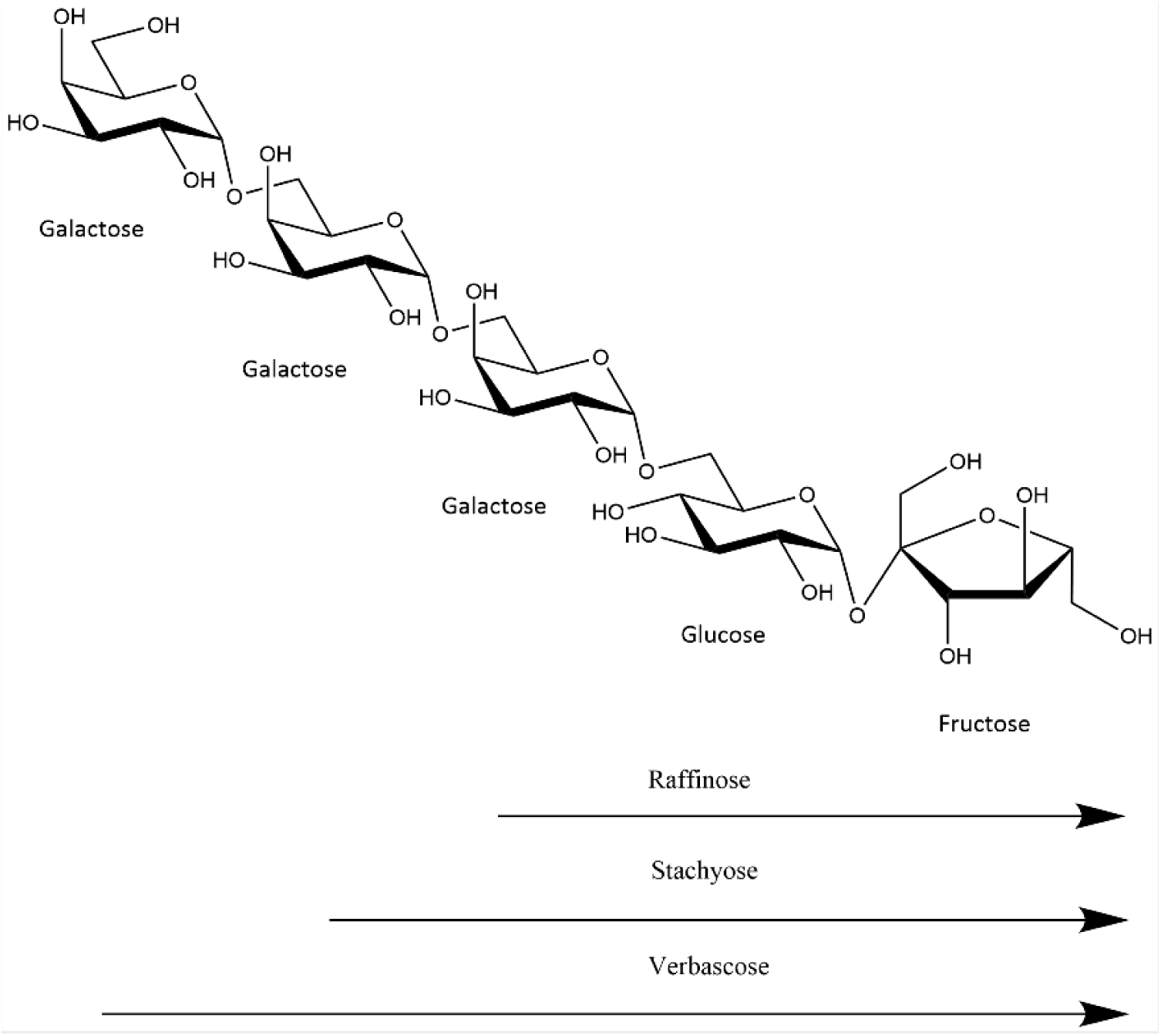
Chemical structure of the raffinose family oligosaccharides

As highlighted above, there are compelling reasons to obtain RFOs from pulses. However, there is a distinct lack of research aiming at extracting RFOs from pulses and even more so beyond the laboratory scale. Thus, this study tested water-based extraction of RFOs with industrially relevant and scalable equipment and the objective of producing a concentrated RFO fraction from commercially available Nordic plant protein materials. Such a process could provide manufacturers with a method to generate an RFO extract that could be further utilized, as well as an RFO-reduced pulse-protein stream while possibly using fewer chemicals as in isolate production.

## 2. Material & Methods

### 2.1. Materials

The field pea (*Pisum sativum L.*) protein concentrate (PPC) was obtained from AM Nutrition (Stavanger, Norway), and the faba bean (*Vicia faba*) protein concentrate (FPC) was purchased from Vestkorn A/S (Tau, Norway). Both are produced by dry fractionation.

Pure water was generated with a Merck Millipore MilliQ system (Burlington, Massachusetts, USA) to reach water with a conductivity of 0.055 µS/cm and 1-2 ppb total organic carbon (TOC). The system was also equipped with an ultraviolet lamp for disinfection.

Analytical standards and all other chemicals, unless otherwise specified, were obtained from VWR (Avantor, Radnor, Pennsylvania, USA) or Sigma-Aldrich (Merck, Darmstadt, Germany).

### 2.2. Biorefinery Process

The biorefining process outlined in Figure 2 was performed in the pilot-scale biorefinery at the Norwegian University of Life Science in Ås, Norway. The process consists of 5 different steps: solubilization in a stirred tank, 2-phase separation, depth filtration of the supernatant followed by ultra- as well as nanofiltration.

**Figure 2:**
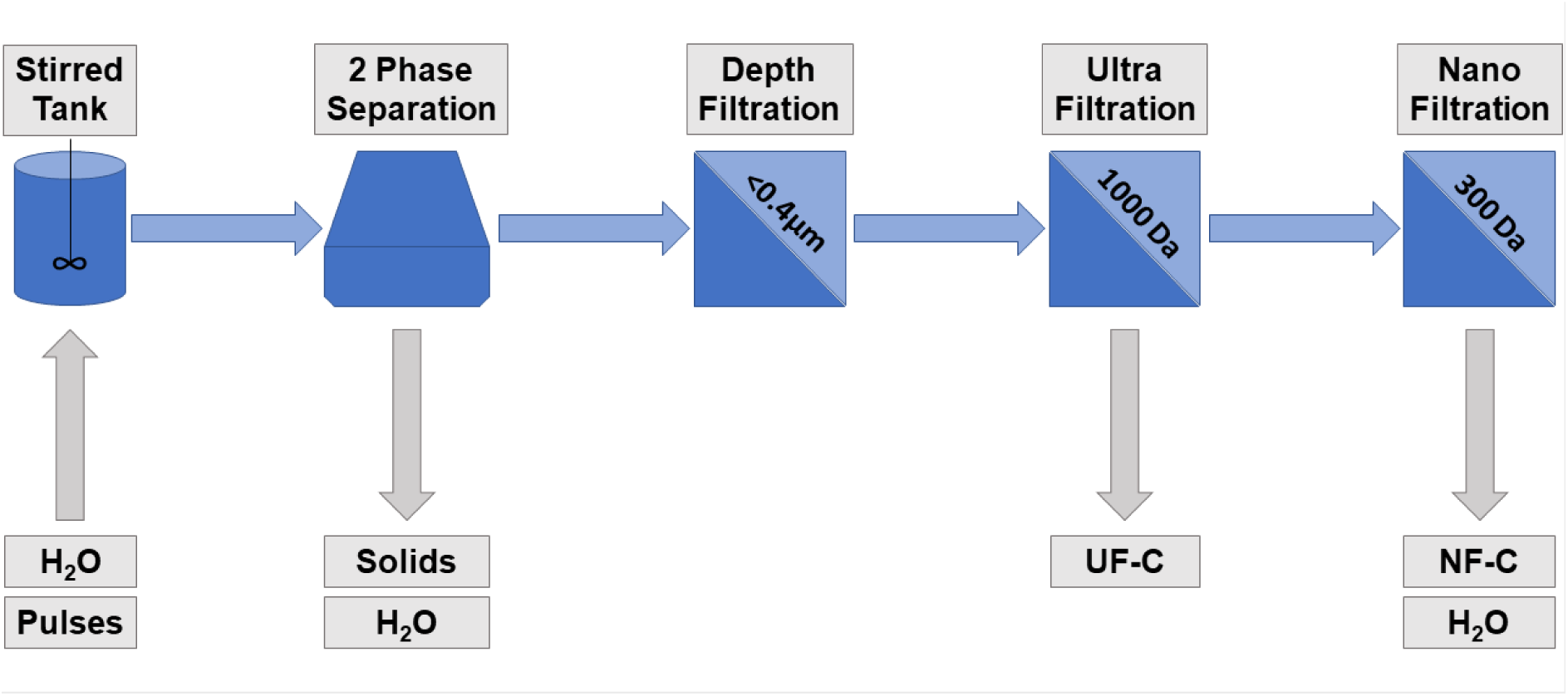
Processing scheme from the solubilization to the final filtration step with the major product and substrate streams (UF-C = Ultrafiltration Retentate, NF-C = Nanofiltration Retentate)

The stirred tank for the solubilization of RFOs had a working volume of 200 L and was equipped with a flat blade impeller (45° angle) stirring at two different heights, a concave floor with a central outlet and temperature control through two separate electrical heating jackets and a thermometer at the bottom of the tank. Solubilization was performed at 40°C and 200 rpm for 1 hour.

The 2-phase disc-stack separator was a GEA Westfalia Easyscale 10.S Unit (Oelde, Germany) operated at 12,000 rpm and 5 bar. The separator was designed to separate liquids of 1.0 kg/dm^3^ from solids of 1.4 kg/dm^3^ and emptying intervals of the separator bowl were adjusted according to manufacturer guidelines based on the suspensions solid matter. Solid matter was determined by centrifuging (4000 rpm, 5 min) two 10 mL samples from solubilization in 15 mL centrifugation tubes and the volume of the pellet was considered as the percentage of insoluble material in suspension.

Depth filtration was performed on a Danmil (Greve, Danmark) 12” DOE Lenticular filter system with 0.2-0.4 µm pore size and total volume of approximately 29 L. The system was filled and washed with pure water before applying material.

Ultrafiltration (UF) and nanofiltration (NF) were performed on a pilot-scale model (GEA Process Enineering A/S, Skanderborg, Danmark) equipped for 65 mm thick (outer diameter) and 965 mm long spiral membranes and a reservoir for retentate/feed. The spiral membranes used in this study were UF: Alfa Laval ETNA 01PP-2538/48 (pore size: 1000 Da) and NF: Alfa Laval NF-2538/48 (pore size: circa 300 Da). Membrane sizes were chosen for all relevant oligosaccharides to pass through UF but being retained during NF. Membranes were stored in 1 % sodium bisulfite and activated before use by 30 min washing in 0.5 mM NaOH (ca. pH 10.5). Afterwards the system was washed with pure water until the permeate had a conductivity of 0 µS cm^-1^. Ultrafiltration was performed at up to 8 bar pressure with a retentate flow of ≥1200 L h^-1^ at a temperature of circa 25-30°C. Nanofiltration was adjusted to 25 bar with otherwise the same conditions. Permeates were collected as soon as the conductivity increased (>0 µS cm^-1^). After applying all material, diafiltration was performed with pure water, holding a relatively constant retentate volume and until permeate conductivity was notably reduced again (see results). After diafiltration, the retentate volume was reduced until the reservoir was empty and then harvested. Harvesting was performed by carefully adding pure water and pushing out the material without mixing back the retentate.

The biorefinery process led to three major streams: 1) A wet solids fraction from the 2-phase separator (“Solids”), 2) an ultrafiltration retentate/concentrate (UF-C) and 3) a nanofiltration retentate/ concentrate (NF-C), as shown in Figure 2. Additionally, there are process-water streams from the separator and nanofiltration step.

Pumping in between the different steps was either performed by the machine’s own pumps or with silicon tubes and peristaltic pumps. If necessary, the processed material was stored in a cold room (4°C) in between process steps. All equipment was rinsed with warm water (up to 80°C) and washed with NaOH solution with pH 10.5 to pH 11 between and after the different steps. The separator and tank were additionally washed with citric acid solution.

The obtained fractions were aliquoted in 500 g samples in aluminum trays, frozen at -20°C and then freeze-dried on a modified freeze-dryer Lyoalfa 15 (Azbil Telstar Technologies S.L.U., Barcelona, Spain) that was constantly held at -80°C and samples were kept in the drying chamber until dry.

### 2.3. Carbohydrate Analysis

The presence of RFOs as well as other carbohydrates in the processing fractions were detected by matrix assisted laser desorption ionization time of flight mass spectrometry (MALDI-ToF MS) on a Bruker ultrafleXtreme (Billerica, Massachusetts, USA). Samples were diluted with pure water and 1 µL was spotted on the target plate with 2 µL 9 mg mL^-1^ 2,5-dihydroxybenzoic acid in 30% acetonitrile and dried before analysis. Masses in the spectrum from 300 to 1000 Da (corresponding to RFOs) were observed during this analysis.

For quantification, RFOs were extracted by adding 9.5 mL ethanol (50 %v/v) to 50 mg sample, and 0.5 mL of melibiose (0.02%) as internal standard. Samples were incubated at 50°C for 1 h under continuous shaking. Samples were then centrifuged at 4000 rpm and an aliquot was diluted 1:20 with water and filtered (0.22 µm). Sample volumes of 25 µL were injected into a High-Performance Anion Exchange Chromatography with Pulsed Amperometric Detection (HPAEC-PAD) as previously described^9^.

Additionally, the starch content in the different fractions was determined in duplicates with the Total Starch Assay Kit (K-TSTA-100A) from Megazyme (Wicklow, Ireland) using the included Rapid Total Starch protocol.

### 2.4. Protein Analysis

The content of proteins in the different materials and processing fractions were analyzed in duplicates according to Method 990.03 of the Association of Official Analytical Chemists (AOAC) with a Dumas N Combustion procedure^29^. The nitrogen to protein conversion factor used was 6.25.

Total amino acid composition of hydrolyzed protein was determined with a 30+ Amino Acid Analyzer (Biochrom Ltd, Cambridge, England) according to Commission Regulation (EC) No 152/2009^30^. Tryptophane was not included in this analysis.

For gel electrophoresis, sample solutions with approximately 2 g L^-1^ protein were prepared and mixed with reducing agent (dithioreitol) and sample buffer (lithium dodecyl sulfate) to a final concentration of 1 g L^-1^ protein. The solutions were boiled for 10 min and then centrifuged, before 10 µL (circa 10 µg protein) were applied to a Mini-PROTEAN TGX stain free gel. The BioRad Precision Plus Protein Standard (unstained) was applied as molecular weight reference. Gels were run at 240 V, 400 mA for 18 min in a BioRad Mini-PROTEAN Tetra System and BioRad PowerPac Basic. Gels were analyzed by exposing the gel for 5 min with a BioRad Gel Doc EZ and band detection was optimized for detecting faint bands. Equipment and reagents were obtained from BioRad (Hercules, CA, United States) and Invitrogen (Waltham, MA, United States) separately.

### 2.5 Ash Content

The ash content was determined in duplicate in accordance with ISO 5984 and the Commission Regulation (EC) No 152/2009 ^30^ by burning all organic material in the sample at 550°C in a furnace (Nabertherm, Germany).

### 2.6. Statistical Analysis

Data was tested for significant differences using analysis of variance (ANOVA) and Tukey’s Honestly Significant Difference test, when there was a significant (*P*<0.05) difference between samples. All statistical analysis was performed in R 4.3.2.

## 3. Results

### 3.1. Method Development

In a first iteration, the process was run without any pH adjustments to reduce the use of chemicals in the process. Pea-protein concentrate had notable flocculation and foam formation during UF which resulted in a drastic decrease in cross-membrane flow. Oligosaccharides with corresponding molecular weight of raffinose, stachyose and verbascose were however detected (MALDI-ToF) in the NF-C fraction. Therefore, the centrifugation/separation behavior of protein concentrates was assessed at different pH values. Protein concentrate suspensions (10 %) with native pH 6.3 were adjusted to different pH from 4 to 7 with 0.5 pH intervals, using 2 M citric acid, then centrifuged (4700 rpm, 20 min) and assessed. After centrifugation (Figure 3), it was observed that adjusting the pH to 5 for PPC and 5.5 for FPC led to a clear supernatant, whereas at pH ≥ 6 centrifugation was not able to separate the suspensions sufficiently. Hence, adjustment to pH 5 with citric acid was chosen to be used in further processing to allow better separation in the 2-phase separator and less reduction of cross-membrane flow during UF.

**Figure 3:**
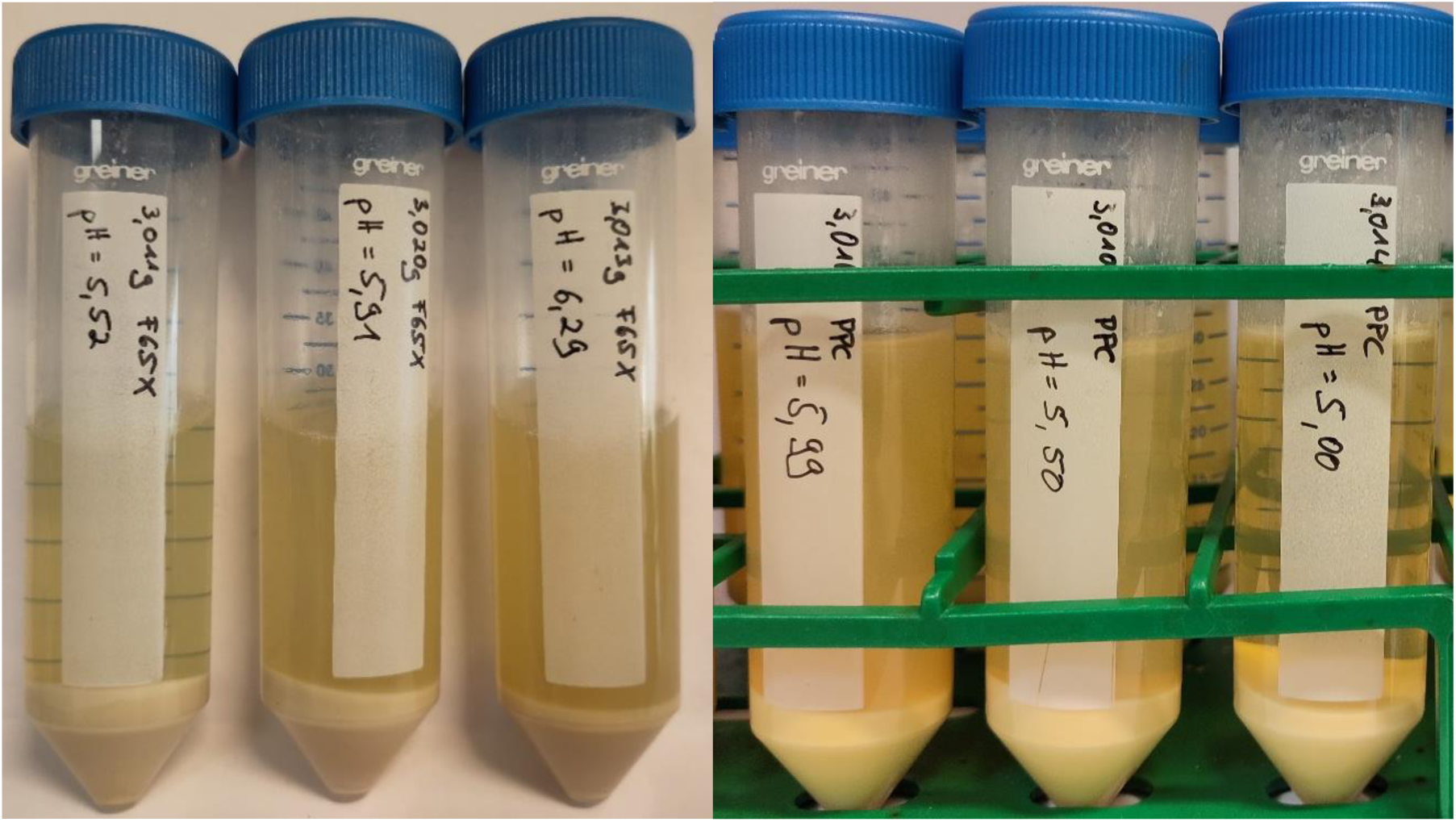
Separation of faba bean protein concentrate FPC (left) and pea protein concentrate PPC (right) suspensions at different pH values after centrifugation

### 3.2. Processing Mass Balances

The obtained masses (wet basis) of all major process streams are shown in Table 1.

**Table 1:** Weight of all major process streams on a wet basis (kilograms) and on a dry basis after lyophilization (grams)

| Step | Solubilization | Separation<br>Solids | Separation<br>Supernatant | Depth<br>Filtration<br>Permeate | Ultrafiltration<br>Retentate<br>(UF-C) | Ultrafiltration<br>Permeate | Nanofiltration<br>Retentate<br>(NF-C) | Nanofiltration<br>Permeate* | Loss* |
| --- | --- | --- | --- | --- | --- | --- | --- | --- | --- |
| PPC – wet [kg] | 110 | 42.5 | 70.9 | 90.0 | 8.7 | 155.2 | 16.0 | 249.2 |  |
| – dry [g] | 10,000 | 8,098 |  |  | 271 |  | 1,186 |  | 446 |
| FPC – wet [kg] | 110 | 73.2 | 66.6 | 106.9 | 6.6 | 170.3 | 11.8 | 326.7 |  |
| – dry [g] | 10,000 | 6,876 |  |  | 197 |  | 978 |  | 1,950 |

\*Calculated values.

#### 3.2.1 Solubilization & Separation

Adjusting of pH during solubilization resulted in the consumption of 1 L and 1.3 L 2 M citric acid for PPC and FPC respectively. In both cases pH stabilized at 5.1. The concentrate suspensions had an approximate solids content of 19.5 % v/v (PPC) and 17 % v/v (FPC) resulting in separator settings of 75 l h^-1^ feed rate with 60 s intervals for bowl emptying. Apart from the resulting streams listed in Table 1, 2-phase separation yielded 60.68 kg (PPC) and 94.1 kg (FPC) process water, which was opaque, light yellow and had a distinct pea / bean odor. Processing left behind some material residues in the stirred tank despite an attempt to flush out residues from the tank and separator with additional pure water. This water increased total mass during the separation stage. For FPC, notable built up of sediment (aka loss of material) was discovered inside the separator bowl after processing.

#### 3.2.2 Depth Filtration

As there is no indicator such as conductivity on the depth filtration unit, product collection was based on a change in color and odor of permeate which appeared after the first 15 kg (half filter volume) of permeate for both materials. The same indication was used when flushing out material with pure water, which resulted in an increase of mass for both fractions, but more so for FPC. Used filters showed a clear accumulation of non-soluble material on the membrane surface.

#### 3.2.3 Ultrafiltration

Circa 3-4 L of permeate passed the membrane before an increase in conductivity was observed and collection was started for both materials. For PPC the conductivity reached >3000 µS cm^-1^ under the used conditions, for FPC only >2500µS cm^-1^ were observed. Using circa 70 kg of pure water reduced conductivity of permeate from PPC to <270 µS cm^-1^, but only to <350 µS cm^-1^ for FPC.

#### 3.2.3 Nanofiltration

As during ultrafiltration, the first few liters of permeate were without conductivity (0 µS cm^-1^). For PPC the conductivity rose to >1400 µS cm^-1^ which could be reduced to <300 µS cm^-1^ by diafiltration with 111 kg of pure water. FPC on the other hand only reached up to 200 µS cm^-1^ which subsequently could be reduced to <80 µS cm^-1^ through diafiltration with 168 kg water. Harvesting of retentate was performed twice to determine the amount of material left in the system due to mixing.

#### 3.2.4 Dried material analysis

After freeze-drying, the dry weights shown in Table 2 were obtained for the different fractions. Compared to 10 kg of used protein concentrates, a loss of roughly 4.5 % dry matter occurs for PPC but 19.5 % for FPC. Furthermore, the initial harvesting of concentrates from the NF system contained most of the dry material (1135 g PPC, 933 g FPC), whereas the second harvesting only contributed with minor amounts of material (51 g PPC, 44 g FPC). All following analytical values for NF-C were a combination of the two harvests. The overall largest fraction on a dry basis (Table 1) during processing were the solids from separation (6-8 kg), followed by the NF-C (circa 1 kg) with the UF-C being the smallest fraction of dry material (0.2-0.3 kg).

**Table 2:**
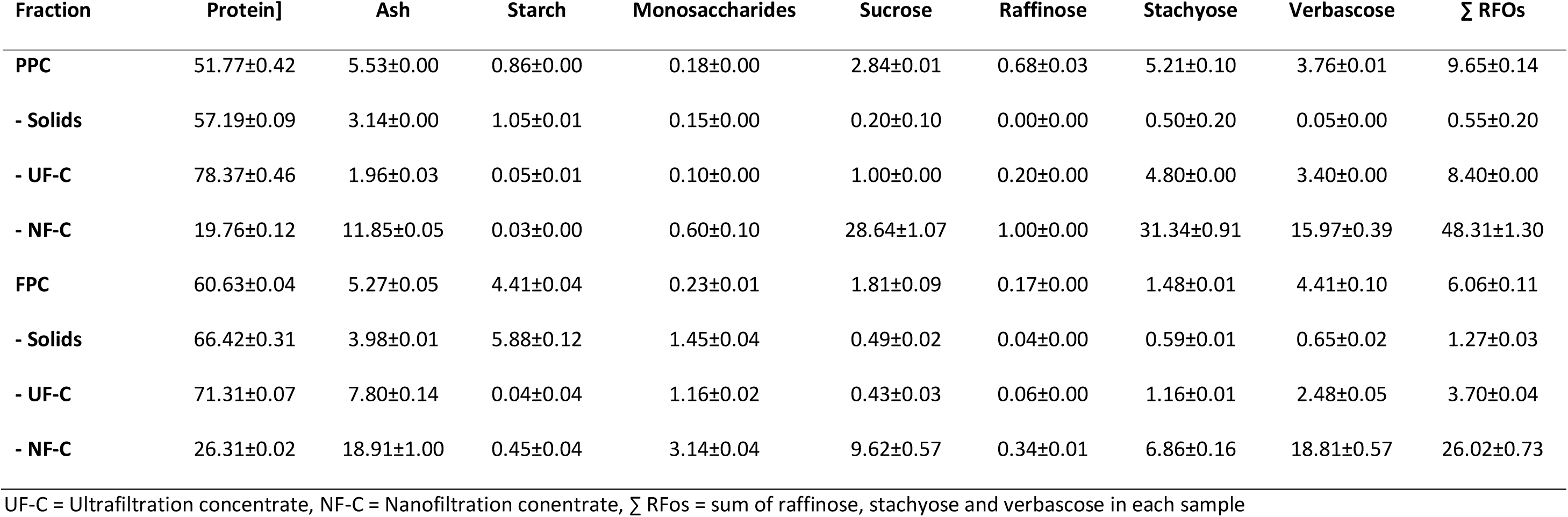
Protein, ash and carbohydrate composition of all obtained fractions on a dry basis [g 100 g^-1^].

### 3.3. Carbohydrate Contents

The process led, as desired, to the production of a fraction (NF-C) that is enriched in carbohydrates, especially RFOs (Figure 4B), and contained significantly more RFOs (*P*<0.001) than all other fractions (Figure 4A). The mass-wise biggest fraction (solids) meanwhile had reduced oligosaccharides content from 9.65 to 0.55 g 100 g^-1^ (-94 %) for PPC and from 6.06 to 1.27 g 100 g^-1^ (-79 %) for FPC compared to their respective substrate (Figure 4A). Not only the reduction was less for FPC, but so was the RFO content in the FPC (Figure 4 A & B) as well as the resulting fractions, especially the NF-C: 26.02 g 100 g^-1^ compared to 48.31 g 100 g^-1^ from PPC. For both materials the UF-C had a carbohydrate content lower than the substrate, but higher than the solids fraction from the separator. Besides RFOs, sucrose was the major carbohydrate in all fractions, with the substrates containing 1.8-2.8 g 100 g^-1^ and the nanofiltration concentrates 9.62-28.64 g 100 g^-1^ for FPC and PPC, respectively. Monosaccharides were only minorly present in all fractions. A detailed overview of all analyzed mono-, di- and oligosaccharides is listed in Table 2.

**Figure 4:**
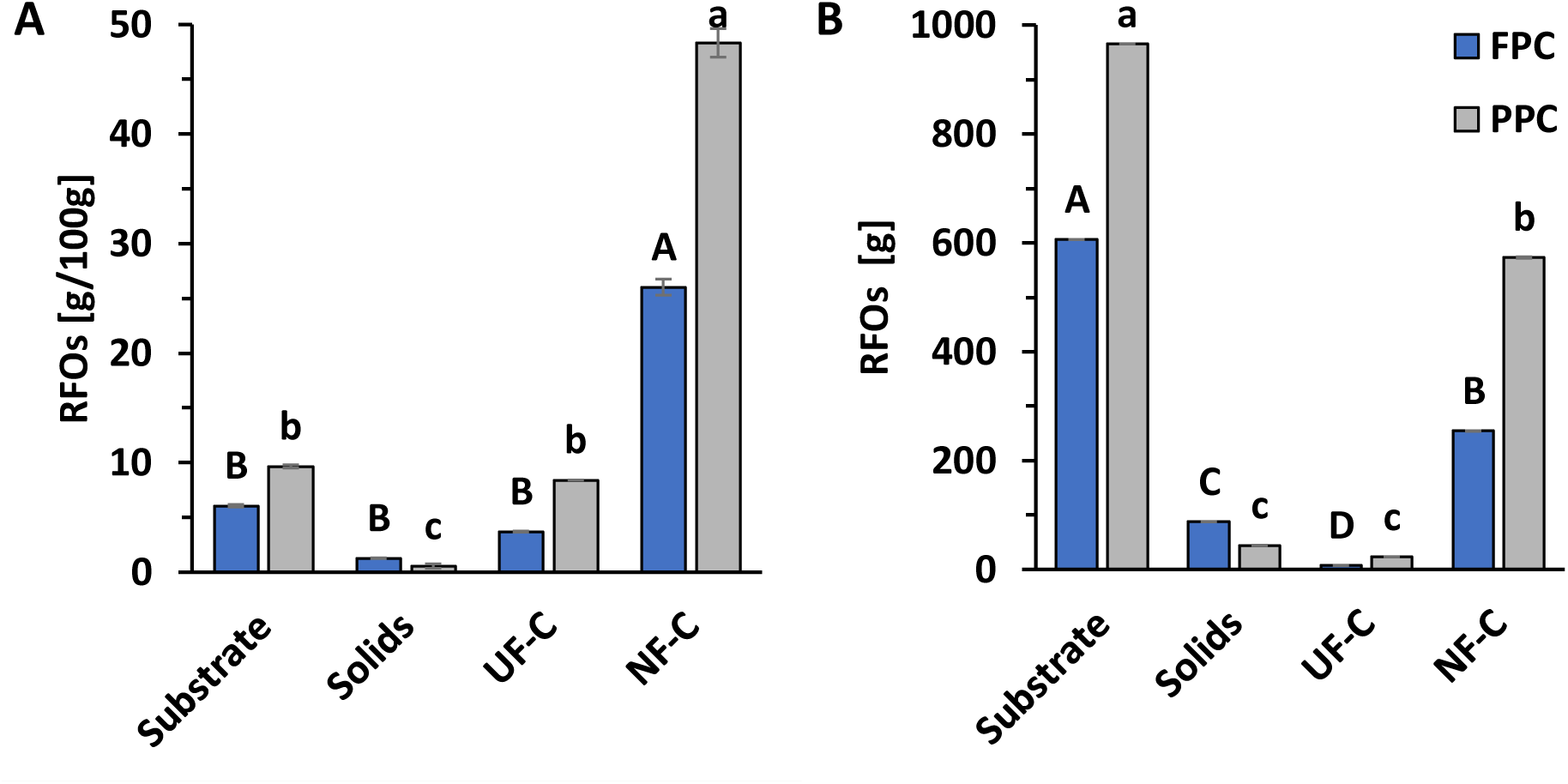
**A)** Concentration and **B)** total mass of raffinose family oligosaccharides (RFOs) in the processing fractions, with UF-C and NF-C denoting ultrafiltration concentrate and nanofiltration concentrate. Error bars indicate standard deviations (n=2). Upper- and lower-case letter denote significant (*P*<0.05) differences among fractions obtained from n faba bean protein concentrate (FPC) or n pea protein concentrate (PPC), respectively.

The starch content in all fractions of pea was ≤ 1 % with the highest content found in the starting material (0.86 g 100 g^-1^) and the solids fraction (1.05 g 100 g^-1^), as shown in Table 2. The faba bean fractions, however, generally contained higher amounts of starch with up to 6.11 g 100 g^-1^ in the solid fractions and 4.41 g 100 g^-1^ in the starting material. All filtration products contained only low amounts of starch (PPC<0.1 & FPC <0.5 g 100 g^-1^).

### 3.4. Protein Contents

The protein content increased from the raw material to the solids fraction, as shown in Figure 5 A. The increase for pea is around 5.42 g 100 g^-1^ and for faba bean around 5.79 g 100 g^-1^. The fractions that remained after ultrafiltration are even more concentrated in protein with 78.37 and 71.31 g 100 g^-1^ for PPC and FPC respectively, but these fractions are much lower in total mass (Table 1) and make up 4.1 and 2.3 % of total protein respectively (Figure 5B). The process effectively kept most proteins from being extracted, but based on the used analysis there was a carry-over of protein into the final carbohydrate concentrate after nanofiltration. For PPC, this is 19.39 and for FPC 26.12 g 100 g^-1^, corresponding to 4.53 and 4.24 % of the total protein respectively and depict a source of protein loss.

**Figure 5:**
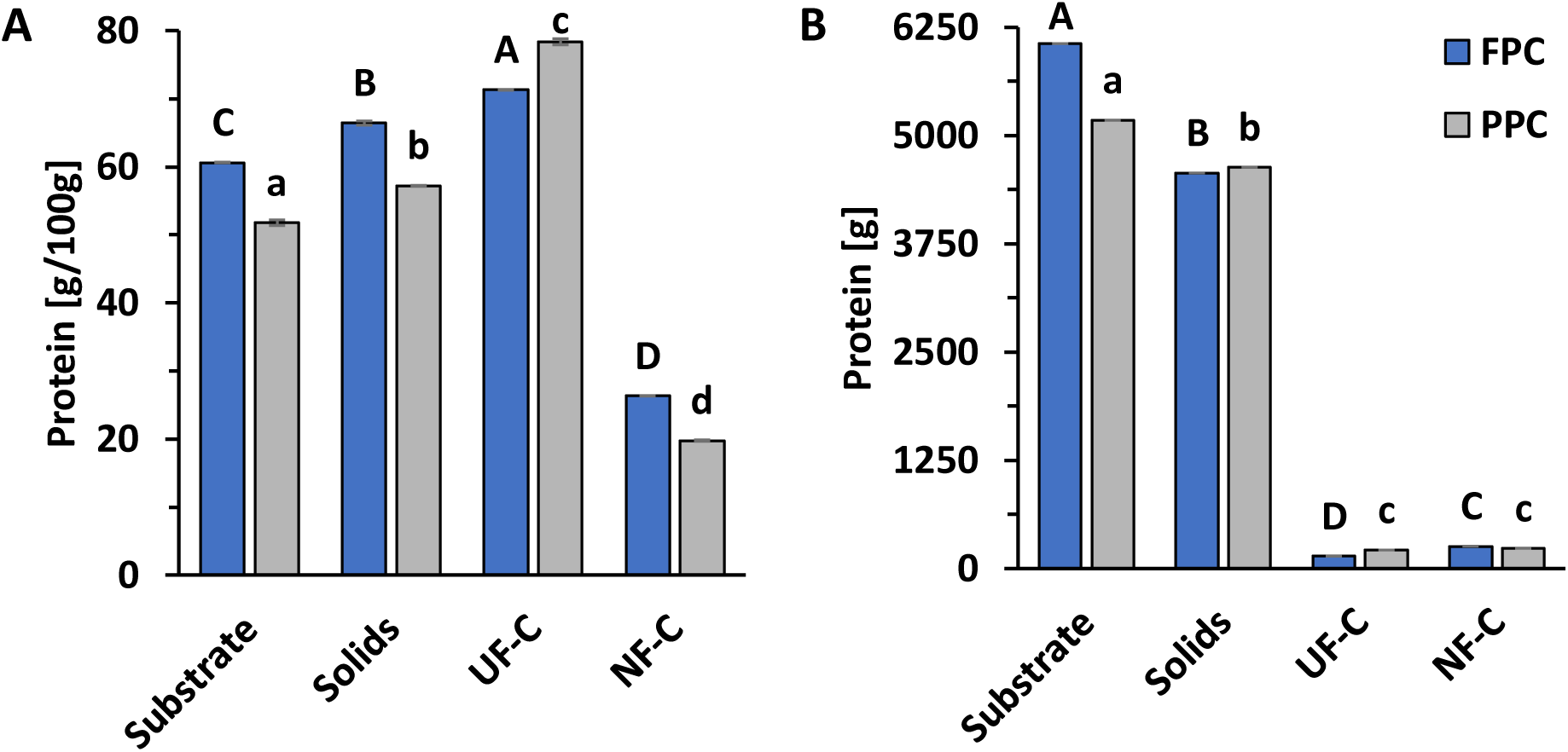
**A)** Concentration and **B)** total amount of protein in the processing fractions, with UF-C and NF-C denoting ultrafiltration concentrate and nanofiltration concentrate. Error bars indicate standard deviations (n=2). Upper- and lower-case letter denote significant (*P*<0.05) differences among fractions obtained from n faba bean protein concentrate (FPC) or n pea protein concentrate (PPC), respectively.

Gel electrophoresis of the protein rich fractions revealed that the solids fraction from PPC (Figure 6) mainly consists of higher molecular weight (MW) proteins (50-100 kDa) whereas the ultrafiltration retentate consisted of lower MW proteins (10-25 kDa), but also higher MW proteins to some degree. The mixture of these two fractions (lanes 9 & 10, Figure 6) resembled the unprocessed PPC fraction (lane 12 & 13 in Figure 6) with major bands around 100, 75, 37, 25, 20 and between 10 & 15 kDa. The unprocessed material had however less accumulation of proteins below 10 kDa. The gel for FPC (not shown) displayed the same trend, a high concentration of lower MW proteins in the ultrafiltration fraction, but also some higher MW proteins and mostly higher MW proteins in the solids fraction. The combination of both fractions showed the same profile as the untreated material. The amino acid (AA) profile (Table 3) of the substrate and solids fraction was very similar to each other, with cysteine and methionine being the least and glutamic acid being the most abundant amino acids. Glutamic acid remained the most and methionine remained the least abundant AAs in the ultrafiltration fraction as well, cysteine however was increased. Generally, the concentration of AAs did increase through processing. The total amount of AAs increased by circa 4 to 5 g 100 g^-1^ from the substrate to the solids fraction for pea and faba bean respectively, but by 27 and 12 g 100 g^-1^ for the UF-C of pea and faba bean respectively. When the protein content of each fraction (Table 2, Figure 5A) is factored in to the sum of the amino acids, the difference in total AAs between the substrate and solids fraction was 1 g 100 g^-1^ or less, and the difference for the substrate and the ultrafiltration product reduced to between 5 and 8 g 100 g^-1^. Of the essential amino acids, leucine and lysine generally had the highest concentrations. However, in the UF-C leucine was the only essential AA that is significantly reduced compared to the starting material, for both pea and faba bean.

**Figure 6:**
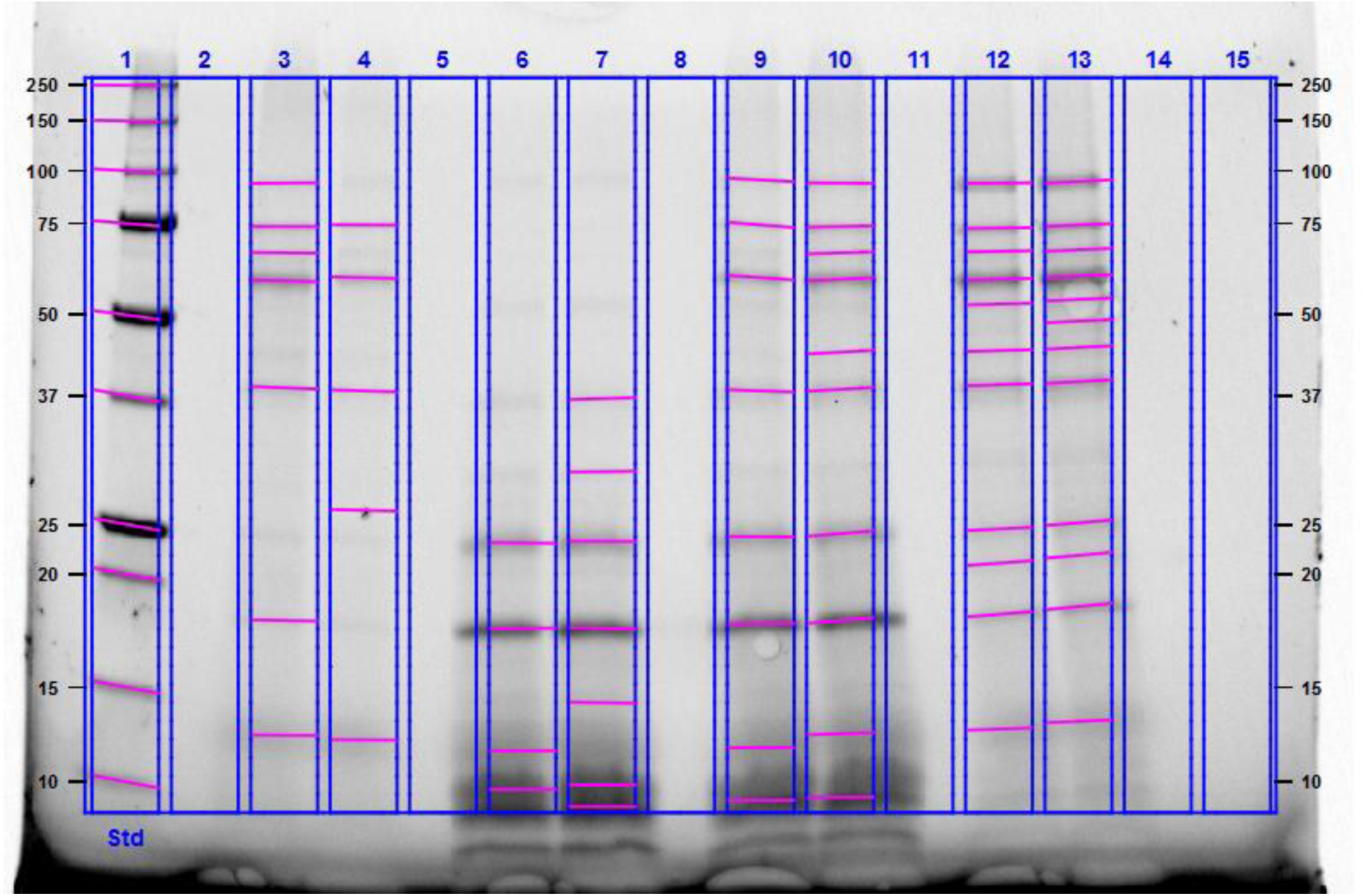
SDS-Page gel of different fractions from pea protein concentrate (PPC). Lane 1 = Standards (Std), lanes 3+4 = Solids Fraction, lanes 6+7 = Ultrafiltration concentrate (UF-C), lanes 9+10 = Mix of Solids and UF-C, lanes 12+13 = unprocessed PPC.

**Table 3:** Content of amino acids of the major protein fractions in the process in g/100g dry matter with standard deviation. Letters in superscript denote significant (p<0.05) differences for each amino acid according to Tukey’s test.

| Fraction | Cys<sup>1</sup> | Met<sup>1</sup> | Asp | Thr<sup>1</sup> | Ser | Glu | Pro | Gly | Ala | Val<sup>1</sup> | Ile<sup>1</sup> | Leu<sup>1</sup> | Tyr<sup>1</sup> | Phe<sup>1</sup> | His<sup>1</sup> | Lys<sup>1</sup> | Arg | Σ | Σ* |
| --- | --- | --- | --- | --- | --- | --- | --- | --- | --- | --- | --- | --- | --- | --- | --- | --- | --- | --- | --- |
| **PPC** | 0.45± | 0.45± | 4.83± | 1.64± | 2.06± | 7.38± | 1.84± | 1.50± | 1.64± | 2.00± | 1.92± | 3.16± | 1.29± | 2.15± | 1.14± | 3.62± | 4.24± | 41.31± | 79.80± |
|  | 0.00<sup>CD</sup> | 0.01<sup>C</sup> | 0.03<sup>E</sup> | 0.00<sup>D</sup> | 0.02<sup>E</sup> | 0.07<sup>F</sup> | 0.03<sup>D</sup> | 0.02<sup>E</sup> | 0.03<sup>E</sup> | 0.05<sup>E</sup> | 0.06<sup>D</sup> | 0.01<sup>C</sup> | 0.05<sup>C</sup> | 0.04<sup>C</sup> | 0.01<sup>F</sup> | 0.04<sup>D</sup> | 0.07<sup>D</sup> | 4.33<sup>F</sup> | 8.36<sup>B</sup> |
| **- Solids** | 0.42± | 0.46± | 5.33± | 1.64± | 2.29± | 8.38± | 2.02± | 1.58± | 1.76± | 2.29± | 2.27± | 3.82± | 1.38± | 2.47± | 1.23± | 3.52± | 4.77± | 45.63± | 79.79± |
|  | 0.00<sup>D</sup> | 0.02<sup>C</sup> | 0.01<sup>D</sup> | 0.04<sup>D</sup> | 0.03<sup>D</sup> | 0.08<sup>E</sup> | 0.01<sup>CD</sup> | 0.01<sup>E</sup> | 0.01<sup>DE</sup> | 0.02<sup>D</sup> | 0.02<sup>C</sup> | 0.02<sup>B</sup> | 0.02<sup>C</sup> | 0.05<sup>B</sup> | 0.00<sup>E</sup> | 0.00<sup>E</sup> | 0.02<sup>C</sup> | 1.45<sup>E</sup> | 2.54<sup>B</sup> |
| **- UF** | 2.01± | 0.79± | 8.65± | 4.12± | 3.88± | 10.66± | 3.49± | 3.10± | 4.08± | 3.05± | 2.71± | 2.69± | 2.74± | 3.38± | 1.94± | 6.96± | 4.30± | 68.56± | 87.49± |
|  | 0.02<sup>B</sup> | 0.03<sup>A</sup> | 0.03<sup>A</sup> | 0.07<sup>A</sup> | 0.00<sup>A</sup> | 0.00<sup>A</sup> | 0.10<sup>A</sup> | 0.01<sup>A</sup> | 0.03<sup>A</sup> | 0.01<sup>A</sup> | 0.02<sup>A</sup> | 0.00<sup>D</sup> | 0.09<sup>A</sup> | 0.01<sup>A</sup> | 0.00<sup>A</sup> | 0.04<sup>A</sup> | 0.06<sup>D</sup> | 1.50<sup>A</sup> | 1.91<sup>A</sup> |
| **FPC** | 0.46± | 0.44± | 5.39± | 1.79± | 2.38± | 9.05± | 2.35± | 1.74± | 1.86± | 2.32± | 2.24± | 3.84± | 1.43± | 2.14± | 1.38± | 3.60± | 5.70± | 48.10± | 79.33± |
|  | 0.00<sup>CD</sup> | 0.01<sup>C</sup> | 0.02<sup>D</sup> | 0.02<sup>D</sup> | 0.06<sup>D</sup> | 0.03<sup>D</sup> | 0.04<sup>BC</sup> | 0.00<sup>D</sup> | 0.00<sup>D</sup> | 0.02<sup>D</sup> | 0.02<sup>C</sup> | 0.03<sup>B</sup> | 0.01<sup>BC</sup> | 0.03<sup>C</sup> | 0.01<sup>D</sup> | 0.01<sup>DE</sup> | 0.08<sup>B</sup> | 0.40<sup>D</sup> | 0.66<sup>B</sup> |
| **- Solids** | 0.47± | 0.46± | 6.01± | 1.96± | 2.75± | 10.18± | 2.57± | 1.90± | 2.02± | 2.62± | 2.52± | 4.47± | 1.60± | 2.44± | 1.54± | 3.83± | 6.05± | 53.38± | 80.37± |
|  | 0.00<sup>C</sup> | 0.02<sup>C</sup> | 0.04<sup>C</sup> | 0.03<sup>C</sup> | 0.04<sup>C</sup> | 0.05<sup>B</sup> | 0.04<sup>B</sup> | 0.00<sup>C</sup> | 0.00<sup>C</sup> | 0.01<sup>B</sup> | 0.02<sup>B</sup> | 0.01<sup>A</sup> | 0.02<sup>B</sup> | 0.04<sup>B</sup> | 0.01<sup>C</sup> | 0.00<sup>C</sup> | 0.05<sup>A</sup> | 0.17<sup>C</sup> | 0.26<sup>B</sup> |
| **- UF** | 2.31± | 0.67± | 7.24± | 3.08± | 3.41± | 9.73± | 3.43± | 2.77± | 3.59± | 2.48± | 2.66± | 2.62± | 2.56± | 2.08± | 1.71± | 5.78± | 4.45± | 60.56± | 84.93± |
|  | 0.01<sup>A</sup> | 0.01<sup>B</sup> | 0.12<sup>B</sup> | 0.05<sup>B</sup> | 0.07<sup>B</sup> | 0.24<sup>C</sup> | 0.18<sup>A</sup> | 0.04<sup>B</sup> | 0.07<sup>B</sup> | 0.04<sup>C</sup> | 0.04<sup>A</sup> | 0.06<sup>D</sup> | 0.04<sup>A</sup> | 0.04<sup>C</sup> | 0.01<sup>B</sup> | 0.01<sup>B</sup> | 0.02<sup>D</sup> | 0.96<sup>B</sup> | 1.34<sup>A</sup> |

<sup>1</sup> (Semi-)essential amino acids; Cys = Cysteine, Met = Methionine, Asp = Aspartic Acid, Thr = Tryptophane, Ser = Serine, Glu = Glutamic Acid, Pro = Proline, Gly = Glycine, Ala = Alanine, Val = Valine, Ile = Isoleucine, Leu = Leucine, Tyr = Tyrosine, Phe = Phenylalanine, His = Histidine, Lys = Lysine, Arg = Arginine, Σ = Sum of all amino acids, Σ\* = Sum in g 100 g<sup>-1</sup> Protein

### 3.5. Ash Contents

The ash content was around 5 % for both starting materials and was decreased in the solids fraction, as shown in Table 2, whereas the RFO extract after nanofiltration contains double (pea) or over triple (faba bean) the amount of ash. For UF-C, the results are varying with the pea fraction having a reduced and the faba bean fraction having a slightly increased content. It seems that ash mostly followed the low molecular weight carbohydrate fraction during processing.

## 4. Discussion

The process described in this study led to the desired extraction of RFOs from both field pea and faba bean protein concentrates. The profile of the extracted oligosaccharides aligned with the distribution of RFOs in the substrate but also with literature. In field peas it is common that stachyose is the most abundant, whereas verbascose is usually most abundant in faba beans; while raffinose is the least abundant in both field peas and faba beans^31,32^. Previous studies of different cultivars, years and regions also reported notable amount of sucrose in the seeds^31–33,34^ as in this study (Table 2). The notable co-extraction of sucrose with RFOs from pulses was also described for other extraction methods in literature^35,36^. The strong similarity to the distribution that can be found in the substrates shows that the process keeps the carbohydrate fractions mostly unaltered. The increase in sucrose towards the final extract (NF-C, Table 2), however, hints at residual enzyme activities during processing which cannot be ruled out as leguminous seeds contain α-galactosidases which become activated during germination^23^. This was previously demonstrated in soybeans^37^ and indicated in yellow peas by increased levels of sucrose and monosaccharide and RFO decrease during germination^38^. Monosaccharides, a product of oligosaccharide degradation, are also slightly elevated in the NF-C compared to other fractions despite the membranes pore-size being big enough that they should be washed out.

Compared to a previously described lab-scale process for extraction and purification of RFOs from pea that utilized 50 % ethanol and chromatography^35^, the here presented process extracted 5.7 g RFOs per 100 g PPC without ethanol, which is ≍33 % higher. The difference could come down to variance in the utilized pea cultivar, cultivation location and year as this can change the oligosaccharide content^33^. However, if scaled-up to the same scale as in this study, the process of Gulewicz et al. (2000) would require a total of 80 L pure ethanol for 10 kg of seed material which. A follow-up study of the method described by Gulewicz et al (2000)^35^ utilized even more ethanol to purify RFOs and remove sucrose ^36^. In that regard, the process described in this current study is advantageous, but does not reach the same level of purity. For faba beans, there is no comparable study in terms of α-galactoside extractions, with or without ethanol.

The most similar established process to the current one is wet fractionation, which is usually associated with high water and chemicals consumption. Berghout et al. (2015)^39^, reported that for production of 1 kg of lupin protein isolates 40 g of NaOH as well as 40 g of HCl were necessary to use. The process described in this present study shows that 1kg of PPC or FPC only utilizes 38.25 g or 49.95 g citric acid, respectively, which is 48% or 62% weight of the chemicals during lupin processing, respectively. Furthermore, this process avoids the use of halogens (no chlorine). Numbers for the process are regardless of the cleaning solutions used, which were also not reported in the lupin study^39^.

As described (Table 2), according to Dumas some protein is carried over to the otherwise mainly carbohydrate containing NF-C and the UF-C contains mainly extracted proteins despite the designated protein separation in the 2-phase separator. That it is often possible to further increase protein removal after an initial separation step, was also shown by the additional protein recovery steps in the patent by Markedal et al. (2016)^14^: Even despite the harsher processing conditions (pH 3.5, 50-80°C) in the patent, it seems like protein is always co-extracted in the non-protein fractions, as it was in the current study. The protein-rich UF-C could however be used to increase protein yields in the process by mixing it with the solids fraction. Considering the different concentrations and masses of these fractions, the protein content of the resulting products would increase by another 0.69 and 0.14 g 100g^-1^ for PPC and FPC respectively compared to the solid’s fractions alone. This accounts for 4.1 % (PPC) and 2.3 % (FPC) of all protein in the process. The re-combination of the UF-C and solids would also reintroduce some RFOs. For PPC this would only increase RFO concentrations only by 0.25 g 100g^-1^ and the total decrease would still be above 90 % in comparison to the starting material. For FPC, the amount of RFOs would increase even less (0.07 g 100g^-1^) and the overall decrease would still be around 78% when combining UF-C and solids.

Besides the co-extraction of protein in UF-C and NF-C, a notable source for protein loss in the process could have been the depth filtration step where non-solubilized material >0.4 µm is held back and this could be particles of starch and protein that are hard to separate in dry fractionation, but also slipped the centrifugal separation due to lower particle densities than 1.4 kg dm^-3^. Additionally, protein is likely lost as a residue in the stirred tank (foaming residues on the tank walls) and during separation for FPC (accumulation of matter in the separators disc-stack).

Aside from assessing extraction efficiencies and loss of protein, it was also evaluated whether the process would alter the protein fraction, therefore gel electrophoresis and AA analysis were performed. The SDS page shown in Figure 6 might be over-representing the actual content of low molecular weight proteins in the ultrafiltration retentate, as this fraction was highly concentrated but would only make up a small fraction in a mixture with the solids fraction. However, a solubilization and partial fractionation of proteins in acidic conditions and during processing cannot be completely ruled out. The untreated material gave the best results in SDS Page in terms of clearly visible bands (despite loading similar concentrations of protein), but this was also observed in the gels that are shown in a previous study on pea protein by Kornet et al. (2020)^42^. In the same study, pea flour and different treated fractions contained vicilin of a size of circa 50 kDa, which is also present in the fractions shown in Figure 6 for the here processed pea-protein. Further bands of vicilin are also present, with molecular weights of circa 37 kDa and 14 kDa. Additionally, the legumin beta subunit at around 22 kDa was detected. While Kornet et al. (2020)^42^ ascribe the fraction at 66 kDa to convicilin, Shevkani et al. (2015)^43^ describe this as a 68-70 kDa fraction, which is similar to the detected bands below 75 kDa that the PPC had in this study. The authors of the same study ^43^ assign the band close to 100 kDa, also observed in Figure 6, to lipoxygenase. For proteins in faba bean, a very similar picture arises: Compared to previous studies^11,44^ proteins of beta-legumin type (circa 22 kDa), convicilin (54 & 65-68 kDa) and viciline (48-51 kDa) and potentially also alpha-legumin (37-4 kDa). For both, pea and faba bean protein fractions, it can thus be concluded that the processing did not influence the protein sizes notably when compared to the literature. The composition of amino acids of the here used faba bean concentrate and its processed fractions are comparable to the concentrate in the previous study by Vogelsang-O’Dwyer et al. (2020)^11^: Their study also found glutamic acid to be the most and cysteine and methionine to be the least abundant AAs in the concentrate, but also in the protein isolate included in their study. They also found tryptophane to be among the least abundant AAs, which was however not determined in this study. Their reported overall content of amino acids (57.85 g 100 g^-1^ dry matter) was however closer to the protein content (64.10 g 100 g^-1^ dry matter) than the amount that was found during this study. For pea, a study by Çabuk et al. (2018)^45^ found a similar concentration of AAs (44.16 g 100 g^-1^) and the same AAs are most (glutamic acid, aspartic acid) and least (cysteine, methionine) abundant in the concentrate as well as the here produced solids fraction from pea. The UF concentrate from PPC, but also FPC, deviates slightly as the level of cysteine is increased during the process. Pea and faba bean fractions were very similar in terms of AA profile. Of the essential amino acids, only leucine was significantly reduced in the UF product, but significantly increased in the solids fraction. When combining both fractions this still would result in a net-increase, hence all essential amino acids are increased in the newly produced protein fraction.

Apart from RFOs and proteins, the amount of ash observed in this study is in line with ash contents for other fractions in literature: Faba bean protein concentrates contain around 5 % ash^11, 41^ and for pea protein concentrates 9.6 %^16^ was measured in one study. The reduction of ash in the protein fraction in fact leads to some of the increase in protein, aside from the removal of RFOs. The presence of ash in the RFO extract on the other hand may be acceptable if this material is used as carbon source for microbial fermentations, as ash does not represent a carbon source. Opposite to the ash, the residual starch in the starting materials is co-fractionated with the proteins during 2-phase separation for both substrates. The starch content in all fractions of FPC was higher than PPC despite the higher protein content. This could be due to dehulling prior to air classification which can improve the protein yield and reduce starch content in the case of peas but not faba bean^40^. Some literature does however indicate that the starch content of PPC (1.7 % ^16^) is lower than the content in FPC (>7.55 %^11,41^).

For further process improvement, a decanter centrifuge has the potential to replace the 2-phase centrifugal separator to decrease water in the protein fraction and get a potentially even better oligosaccharide extract and protein enriched fraction. Additionally, it does not need the continuous supply of process water to operate the emptying of the bowl. This could increase processing sustainability and material loss as well (e.g., slight coloring and odor of the processing water from the separator). Generally, a more continuous processing and increased batches might reduce the loss per kg of material, which is inherent in pilot-scale batch processes. The different performances in terms of RFO extraction of pea and bean concentrates indicate the necessity to adapt the process to the different available plant materials. Instead of the equipment, the loading of material could be adjusted, for example based on protein content instead of dry matter. This might be a possibility to avoid fouling in the 2-phase separator, as it was observed during the faba bean processing with higher proteins contents. At the same time, it would also increase the amount of water per gram oligosaccharides and could increase efficiency of the RFO removal.

## 5. Conclusion

By utilizing state of the art process equipment and commercial pulse protein ingredients, a novel biorefining approach to obtain an extract enriched with RFOs could be demonstrated, yielding multiple hundred grams of RFO extract. At the same time, the process effectively decreased the raffinose content in protein concentrates (>75 % removal) and increase protein concentration (>5 % increase) without majorly altering the protein sizes and amino acid composition. Therefore, the process presents a suitable alternative to wet fractionation for ingredient manufacturers, creating a valuable side stream. Upcoming studies will demonstrate the application potential of the RFO extract.

## Acknowledgements

The authors would like to thank Lars Fredrik Moen for all the help with the biorefinery equipment, Elise Hatch Fure for the help with freeze drying and Hanne Hustoft for help with the ash & amino acid analysis.

## Contributions

PG – Main author and responsible for all experiments; SG – HPAEC & protein analysis, methods section, manuscript revision, discussions; BW – Conceiving the study/project, MALDI-ToF analysis, discussions, manuscript revisions; SK – Conceiving the study/project, discussions, manuscript revision; CT – Manuscript revision, study discussion; SS – Manuscript revision, Study discussions.

## Abbreviations

AOAC: Association of Official Analytical Chemists
FODMAP: Fermentable oligo-, oi-, and monosaccharides and polyols
FPC: Fab bean protein concentrate
HPAEC-PAD: High performance anion exchange chromatography - pulsed amperometric detection
LAB: Lactic acid bacteria
MALDI- ToF MS: Matrix assisted laser desorption ionization – time of flight mass spectrometry
MW: Molecular weight
NF: Nanofiltration
NF-C: Nanofiltration Concentrate
PPC: Pea protein concentrate
RFO: Raffinose family oligosaccharides
TOC: Total organic carbon
UF: Ultrafiltration
UF-C: Ultrafiltration Concentrate

